# Maturation-Dependent Changes in the Structure and Seeding Capacity of Aβ42 Amyloid Fibrils

**DOI:** 10.1101/2023.04.11.536374

**Authors:** Alyssa Miller, Sean Chia, Ewa Klimont, Tuomas P.J. Knowles, Michele Vendruscolo, Francesco Simone Ruggeri

## Abstract

Many proteins self-assemble to form amyloid fibrils, which are highly organized structures stabilized by a characteristic cross-β network of hydrogen bonds. This process underlies a variety of human diseases, and can be exploited to develop versatile functional biomaterials. Thus, amyloid aggregation has been widely studied, shedding light on the properties of fibrils and their intermediates. A question that remains open concerns the microscopic processes that underlie the long-time behaviour of the fibrillar assemblies. Here, we use atomic force microscopy to observe that the fibrils undergo a maturation process, with an increase in both fibril length and thickness, and a change in the cross-β sheet content. These changes affect the ability of the fibrils to catalyse the formation of new aggregates through secondary nucleation. The identification of these changes helps us understand the fibril maturation processes, facilitate the targeting of amyloid fibrils in drug discovery, and offer insight into the development of biocompatible and sustainable protein based materials.

## Introduction

The study of protein aggregation and amyloid formation has gained attention over the past few decades because of its association with human disease, such as in the case of the amyloid-β peptide (Aβ) in Alzheimer’s disease (AD). The process of supramolecular aggregation involves the conversion of soluble monomers into large, insoluble, cross-β amyloid fibrils, which possess an extended, highly ordered network of intermolecular hydrogen bonding^1^. When monitored *in vitro* using assays that report on the mass of aggregates structures present, the process of aggregation characteristically involves three phases: (1) a lag phase, where initial aggregates form but their numbers are below the detection threshold^2^, (2) a growth phase, where the aggregates proliferate rapidly, and (3) a plateau phase, where the bulk aggregate mass stops increasing because all the supersaturated monomers have been used up^3^.

At a single-molecule scale, this macroscopic behaviour can be explained by a complex network of microscopic processes. These processes have been investigated through the development of reproducible aggregation reactions in vitro, which can be monitored via fluorescent probes for cross β-sheet structures, such as thioflavin T (ThT), and the development of analytical methods to extract kinetic information from bulk data^4–7^. These analytical approaches have provided key mechanistic insights into the fast processes of aggregation, revealing that the most important microscopic processes, at least during the lag and growth phases, include: (1) primary nucleation, where initial fibril seeds are formed, (2) elongation, where the seeds grow into fibrils, (3) secondary nucleation, where existing fibrils catalyse the formation of new seeds, and (4) fragmentation, where fibrils break to generate new growing ends. Since secondary nucleation and fragmentation of mature fibrillar aggregates, collectively referred to as secondary processes, are mainly responsible for the proliferation of the aggregates, it is important to study them in detail^6^.

The plateau phase can provide a wealth of information about the role of fibrils in secondary processes. However, it has been challenging to obtain mechanistic information about the plateau phase when using standard fluorescent probes. Bulk techniques, such as ThT-fluorescence, provide useful information on the average properties of fibrils in solution. However, their use does not readily capture the subtle changes in heterogeneity characteristic of protein self-assembly processes. Thus, this approach leaves a gap between the *in vitro* study of fibril assembly and the formation of supramolecular fibril assemblies that make up amyloid plaques and inclusions in disease tissues.

Traditional structural methods, such as cryo-electron microscopy (cryo-EM) and nuclear magnetic resonance (NMR) spectroscopy, have provided atomistic descriptions of the core structure of amyloid fibrils ^8–10^. However, it can be difficult to extend these techniques to capture the full breadth of heterogeneity of amyloid fibrils due to averaging effects. Therefore, to investigate fibrils in the plateau phase, one can exploit both bulk methods and single-molecule methods to obtain a more complete picture of the time evolution of the aggregates. Atomic force microscopy (AFM) is a single molecule technique that can be used to measure the time-resolved 3D structural properties of amyloid assemblies at the single-fibril level^11^. However, preparing studies for complementary bulk techniques can be challenging due to the different conditions required by these experiments. For example, bulk aggregation reactions carried out in plate readers are carried out with much lower concentrations of proteins than needed for a bulk infrared (IR) spectroscopy measurements. This makes it impractical to perform parallel studies of multiple fibrillar properties.

Here, we address the problem of identifying the microscopic steps that take place during the plateau phase. Our results illustrate the changes in morphology and structure of heterogeneous, mature Aβ42 fibrils in this phase. We report on the properties of fibrillar aggregates of Aβ42 as a function of time during the early hours of the plateau phase, where ThT fluorescence intensity remains constant. We analyse *in vitro* aggregation experiments by using a combination of single-molecule AFM imaging, Fourier transform infrared (FTIR) spectroscopy, chemical kinetics (**Figure 1**), and a microfluidic deposition^12–14^. Using bulk and single-molecule approaches, we characterize changes in the fibrillar structures during the plateau phase. This information enables investigations of both seeding and secondary processes, which facilitate the development of compounds targeting amyloid aggregates for pharmacological interventions, as well as of protein-based functional materials.

**Figure 1.**
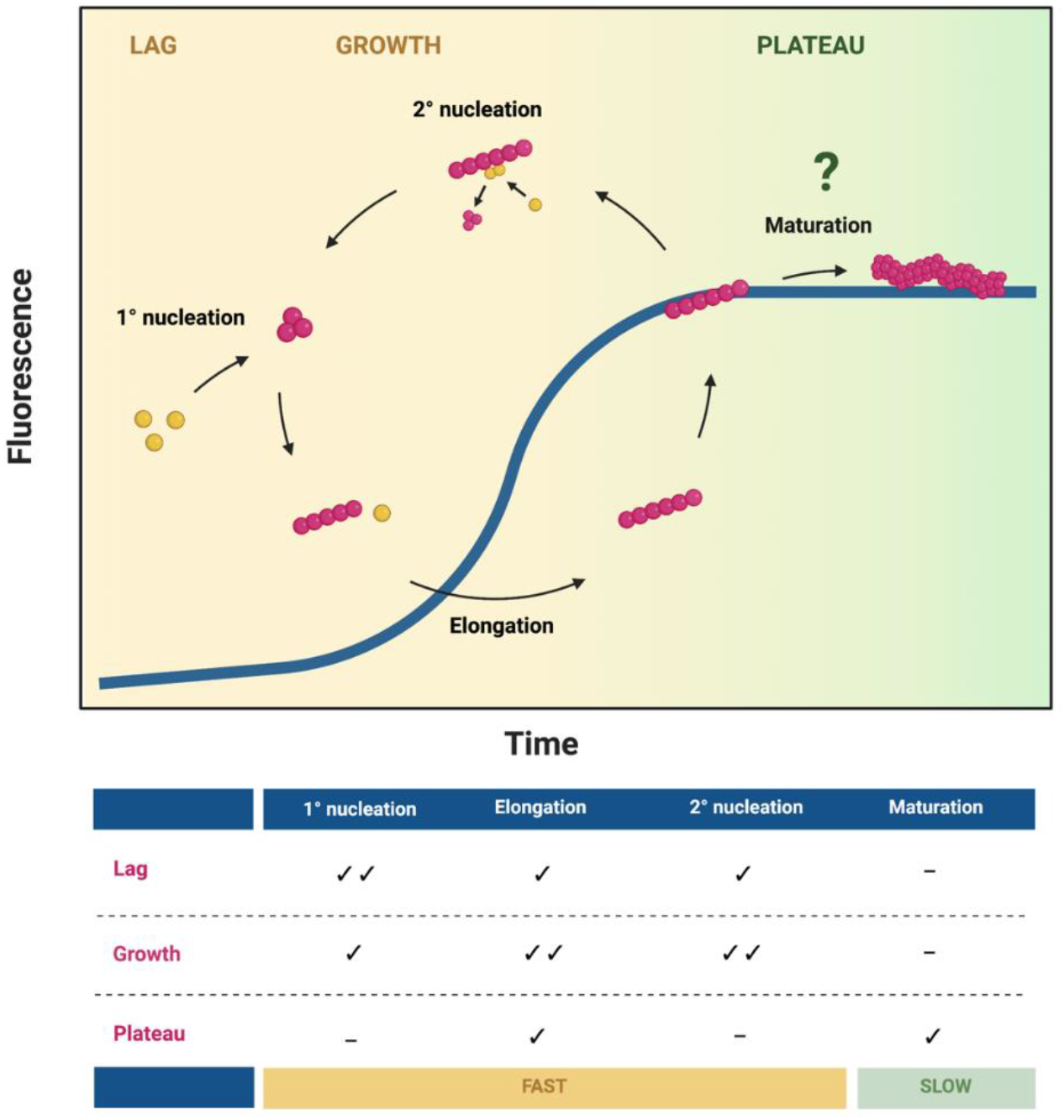
Unravelling Aβ42 fibrils structural changes within the plateau phase. The aggregation of Aβ42 was monitored using a ThT-fluorescence assay, which monitors cross-β structure formation, with a schematic of the dominant molecular mechanisms involved in the aggregation process. Chemical kinetics can shed light on fast processes, however fluorescent probes are insensitive to changes in fibril properties in the plateau phase. Therefore, we characterise slow maturation processes in the plateau phase by studying dynamic structural features of fibrils by AFM and FTIR.

## Results and Discussion

To characterise the properties of fibrillar aggregates in the plateau phase, we first monitored the aggregation of Aβ42 monomers using a ThT-kinetics based assay (**Figure 2**). We defined the beginning of the plateau phase as the region where the growth curve approached a slope of zero. We first sought to assess the ability of fibrils to act as effective templates for conversion of monomers into aggregates, i.e. the seeding capacity, as a function of time. We thus took aliquots from an ongoing reaction at the start of the plateau phase (0 h, early), and at a few hours later (4 h, late), and used these aliquots to generate fibril seeds. These seeds were then reintroduced to monomer solutions, where they acted as templates for self-assembly (**Figure 2**).

**Figure 2.**
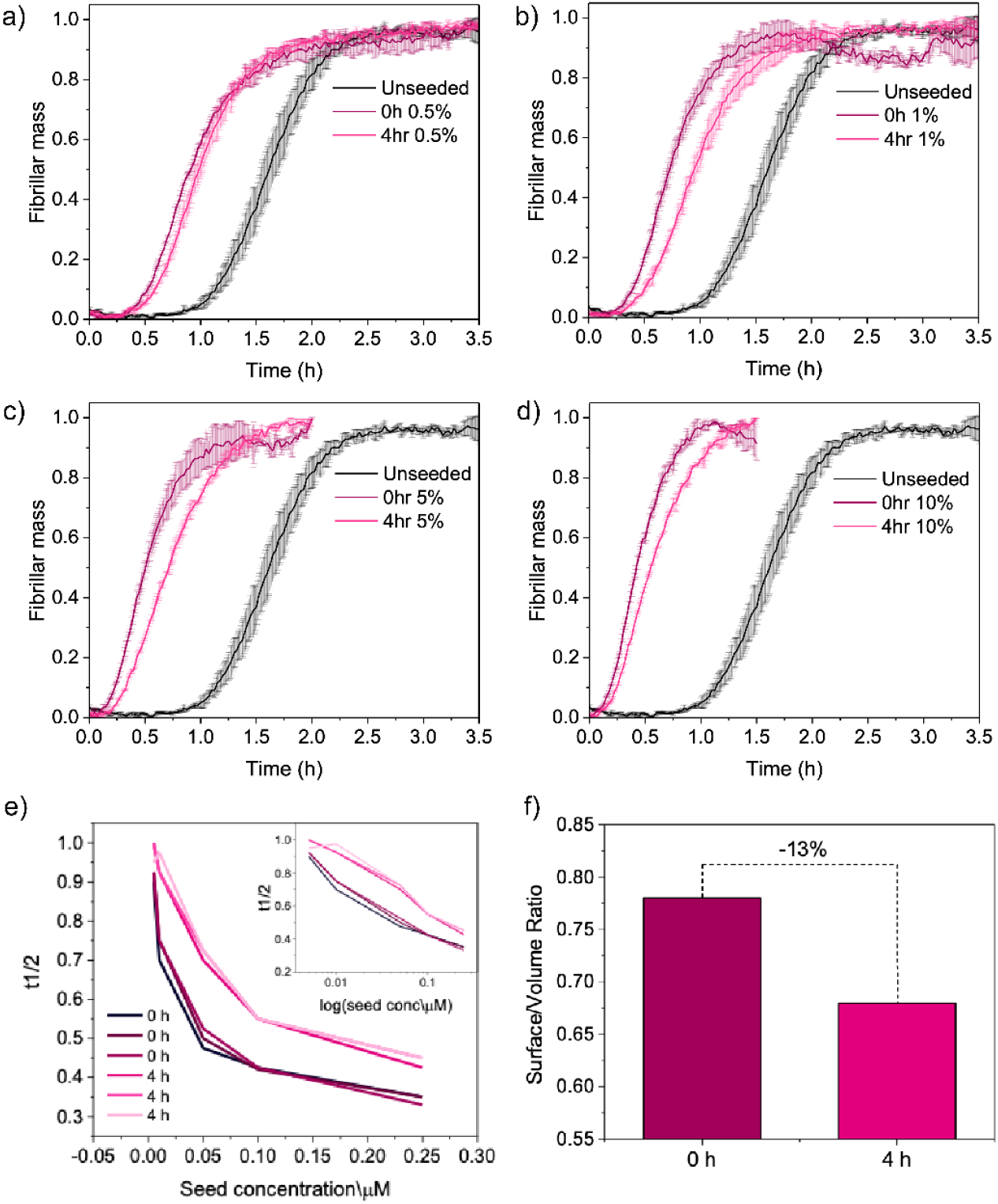
Seeding capacity of Aβ42 fibrils from different time-points in the plateau phase. (a-d) Aβ42 seeds were generated from fibrils in the plateau phase at 0 and 4 h, and the kinetic profiles (average and standard deviation, n=3) were compared to unseeded aggregation. Seed concentrations of 0.5% (a), 1% (b), 5% (c) and 10% (d) were assessed. **(e)** The half-time of the aggregation reaction was plotted as a function of seed concentration. The inset shows the data plotted on a logarithmic scale. Seeds generated from fibrils generated later in the plateau phase exhibit a decreased seeding capacity of 20 ± 10%. **(f)** The surface-to-volume ratio as measured by single fibril imaging decreases by ca. 15% later time points in the plateau phase.

We found that seeds generated from both early and late fibrils were seeding-competent, promoting conversion of monomers into on-pathway amyloid aggregates. At low concentrations of seed (0.5%, **Figure 2a**), where we expect secondary-nucleation to be the dominant mechanism^15^, there was little difference between the early and late seeds. When the seed concentrations were increased to a range where we expect the dominant mechanism to shift to elongation^15^ (1-10%, **Figure 2b-d**), we observed a difference in seeding ability between the fibrils from early and late time points in the plateau phase. In particular, we observed that early seeds acted as more efficient templates for aggregation, speeding up the kinetics of aggregation. To quantify this increase in seeding capability, we measured the half-time (t1/2) of the aggregation reaction. In the conditions studied here, aggregation reactions with early seeds had half times approximately 10-20% shorter than those with late seeds (**Figure 2e**).

To investigate the structural basis for this change in seeding capacity, we performed single-molecule AFM studies. This approach enables the characterisation of the morphological changes at the single fibril-level (**Figure 2f, Figure 3** and **Figure S1**). We took aliquots from an ongoing aggregation reaction at early (0 h) and late (4 h) time points within the plateau phase, as above, and immediately deposited the sample onto a mica surface (**3a**,**b**). We thus acquired phase-controlled maps of the 3D morphology changes of the fibrillar aggregates between the early and late time points of the plateau phase. We could directly measure and quantify an increase in the length, cross-sectional diameter and surface to volume ratio of the fibrils as a function of time in the plateau phase (**Figure 3e,f**) by performing a single molecule statistical analysis. Between early and late time points, the average length of fibrillar aggregates increased from 250 ± 25 nm to 330 ± 25 nm. We also measured the average cross-sectional height of fibrillar aggregates, which increased from 5.15 ± 0.25 nm to 5.90 ± 0.30 nm.

**Figure 3.**
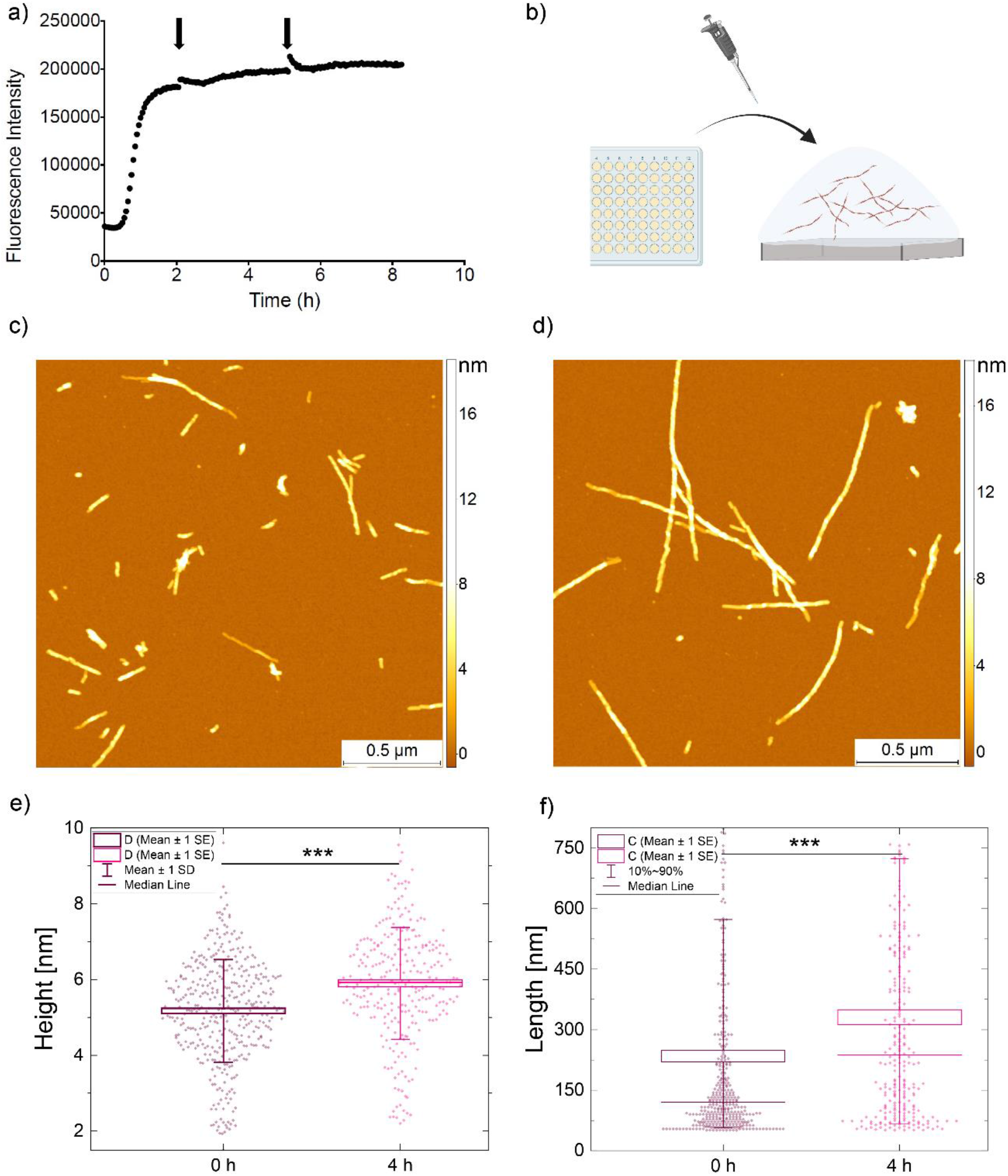
The morphology of Aβ42 fibrils change over time in the plateau phase. (**a**,**b)** Aβ42 fibril formation was monitored using ThT fluorescence, with samples taken from ongoing aggregation reactions (indicated by black arrows) in a 96-well plate, before being deposited on a mica surface. **(c**,**d)** Aβ42 fibrils were imaged via AFM at the start of the plateau phase (c) and 4 h later (d). **(e**,**f)** The height (e) and length (f) of the aggregates were measured for these time points and found to increase over time. Boxes represent ±1 standard error of the mean, lines represent the median, and the whiskers represent the standard deviation. n=389 and 271 for 0 and 4 h, respectively.

Overall, the increase of length of the fibrils is consistent with the continual addition of monomers to the fibril ends over time in the plateau phase. In addition, the increase in the cross-sectional diameter (thickness) of fibrils over time in the plateau phase can be attributed to a hierarchical fibril maturation process, in which single protofibril strands and mature fibrils laterally associate with each other to form higher order structures, resulting in a maturation of the cross-β structure^8,16,17^.

Structural and chemical changes have been reported during the progression from the protofibril to the mature fibril level for a variety of protein systems, which can be affected by aggregation-enhancing mutations.^18–23^ However, detailed characterisations of these changes are still required^24,25^. Therefore, we next sought to address if the seeding capability and morphological changes of the fibrillar aggregates is associated with changes in the secondary structure, as measured via FTIR.

To facilitate comparisons between aggregates measured via AFM, we performed FTIR measurements in the same conditions (**Figure 4**), namely protein concentration (<5 μM) and buffer environment (20 mM sodium phosphate). These conditions typically represent a problem for FTIR measurements in attenuated total reflection (ATR), as the presence of salt crystals causes spectral distortion for samples with lower protein concentration, preventing us from performing spectral analysis in relevant conditions^14^.

**Figure 4.**
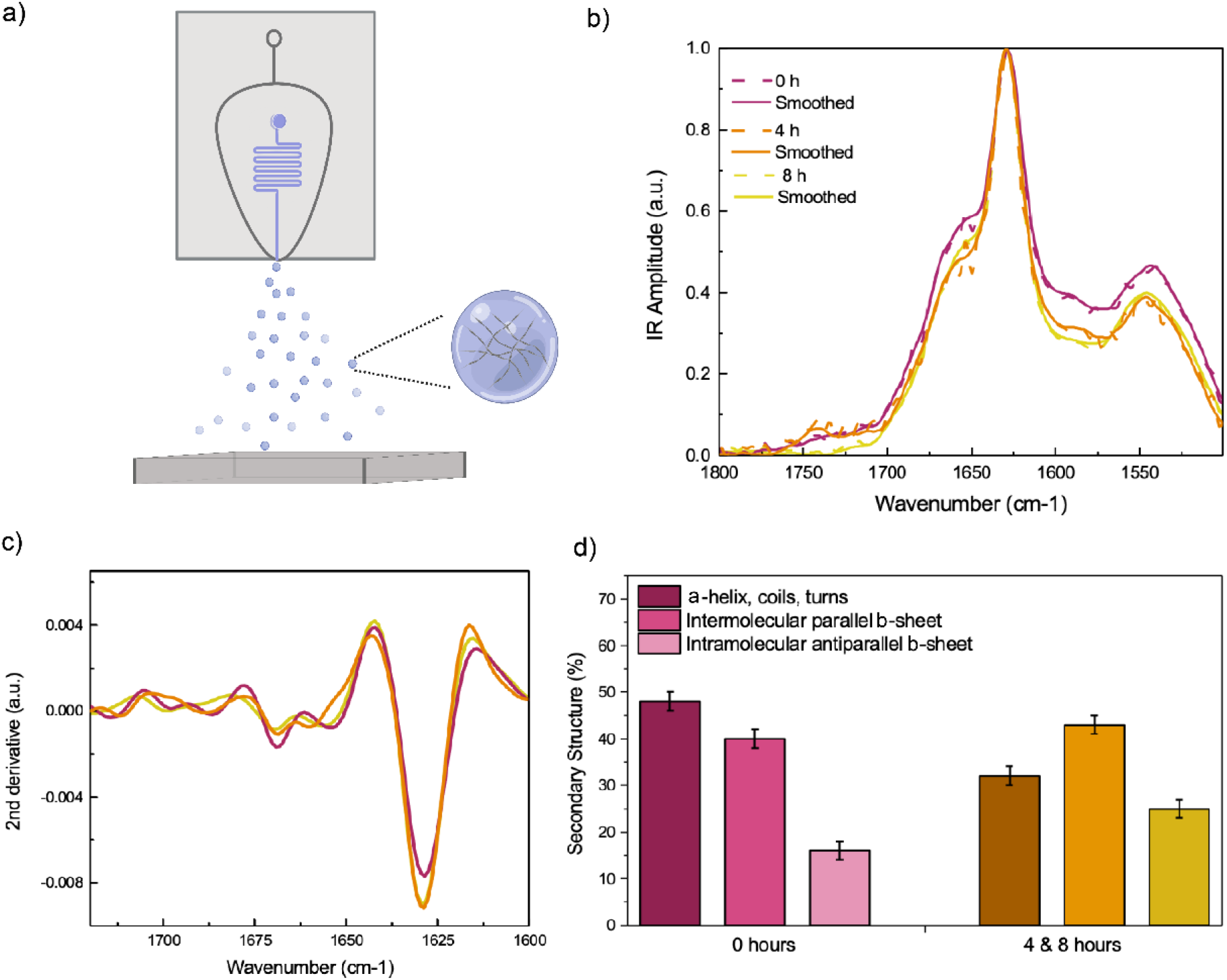
The secondary structure of Aβ42 fibrils, as monitored by FTIR, indicates a net increase in cross-β structure over time. **(a)** A schematic of the microfluidic spray sample deposition onto the FTIR prism. **(b)** Fibrils were taken at 0, 4, and 8 h into the plateau phase and deposited directly onto the ATR prism. Infrared (IR) spectra were acquired for all times points. **(c)** The second derivative was taken for the spectra, revealing a major absorption peak at 1628 cm^-1^, attributed to intermolecular β-sheet. The cross-β signature is more prominent at 4 and 8 h. **(d)** Secondary structure components are shown for fibrils at 0, 4, and 8 h in the plateau phase.

As such, we deposited the sample onto the FTIR prism via microfluidic spray deposition (**Figure 4a** and **Figure S2**)^12–14^. This method minimises salt crystallisation, via ultra-fast droplet drying, to improve the spectroscopic sensitivity such that we were able to acquire spectra with only ∼8.3 ng deposited on the prism (**Figure S3**). This enabled us to measure IR spectra at relevant buffer and concentration conditions (**Figure 4b**), without needing to concentrate the sample, which may mask subtle changes between samples at different time points measured in bulk, while also maintaining the full heterogeneity of the sample ^14^. This also allowed us to obtain measurements consistent to ThT fluorescence and AFM experiments. Then, we used the information in the IR spectra, we assessed the secondary and quaternary structure of the protein aggregates via the analysis of their amide I peak, which contains information on the type of protein fold present^26^. Using this approach, we measured the secondary structure of fibrillar aggregates taken from an aggregation reaction at early (0 h) and later (4 and 8 h) time points into the plateau phase. At both time points, the spectra showed the characteristic signature of amyloid fibrils cross-β sheet structure at 1628 cm^-1^ (parallel) and 1695 cm^-1^ (antiparallel) (**Figure 4c)**. At the early time point, the spectra showed a total cross-β sheet structure of 55±2%, while at the late time points, there is a net increase in this cross-β signature to 69±2% (**Figure 4d)**, as calculated by band integration and the second derivative analysis of the amide I peak^26^.

The changes in secondary and quaternary structure as a function of the time in the plateau phase, and in particular the increase in cross-β structure content, is consistent with an increase in the density of intermolecular hydrogen-bonding network present between β-strands in the fibril ^23,27^. This indicates that the change in cross-sectional diameter is associated with a change in the chemical environment, and therefore the tertiary/quaternary structure of the fibrils under investigation. Such changes in tertiary/quaternary structure of fibrils are associated to a decrease in their surface to volume ratio and reduced seeding capability. As in the case of Aβ42 the fibril surface, as opposed to the fibril ends, acts as the template for seeding^6,28^, this change in fibril structure may explain the variability in seeding capacity described above. The slight decrease in seeding capacity is consistent with the decrease in available fibril surface area to act as a template for misfolding and aggregation, or perhaps further fibril-fibril association masks specific nucleation sites on their surface.

## Conclusions

We have characterised the chemical and structural properties of fibrillar aggregates of Aβ42 within the plateau phase in *in vitro* aggregation reactions. We have identified that within this phase, fibrils undergo a maturation process that includes an increase in cross-sectional diameter (thickness) and length, as well as a change in the fibril hydrogen-bonding network and secondary structure. Such changes in morphology and secondary structure are associated with a variable seeding capacity.

These results shed light on the dynamic processes occurring in the plateau phase of aggregation, contributing to our understanding of the behaviour of mature fibrils. A better understanding of the microscopic processes that take place in the plateau phase increase our knowledge of the protein aggregation process, and help the development of procedures to handle amyloid fibrils for diverse applications, from drug discovery to materials science^29,30^.

## Methods

### Aβ42 production

Expression and purification of the recombinant Aβ (M1-42) peptide (MDAEFRHDSGYEVHHQKLVFFAEDVGSNKGAIIGLMVGGVVIA), denoted Aβ42, were carried out as previously described^31^. Aβ42 was expressed in the *Escherichia coli* BL21Gold (DE3) strain (Stratagene, La Jolla, CA) and purified by sonication and dissolving the inclusion bodies in 8 M urea, followed by ion exchange in batch mode on diethylaminoethyl cellulose resin. Fractions containing Aβ42 were lyophilised and further purified using a Superdex 75 HR 26/60 column (GE Healthcare, Chicago, IL), and eluates were analyzed using SDS−polyacrylamide gel electrophoresis for the presence of the desired protein product. The fractions containing the recombinant peptides were combined, frozen using liquid nitrogen, and lyophilised in 20 mM sodium phosphate buffer, pH 8, 0.2 mM EDTA. The lyophilised products were then stored at −80 °C.

### Chemical kinetics and seeding assays

Solutions of monomeric peptides were prepared by dissolving the lyophilized Aβ42 peptide in 6 M GuHCl. Monomeric forms were purified from potential oligomeric species and salt using a Superdex 75 10/300 GL column (GE Healthcare) at a flow rate of 0.5 mL·min^−1^, and were eluted in 20 mM sodium phosphate buffer (pH 8) supplemented with 200 μM EDTA. The center of the peak was collected, and the peptide concentration was determined from the absorbance of the integrated peak area using ε280 = 1,490 L·mol^−1^ cm^−1^. The obtained monomer was diluted with buffer to the desired concentration and supplemented with 20 μM ThT from a 1 mM stock. All samples were prepared in low-binding Eppendorf tubes on ice using careful pipetting to avoid introduction of air bubbles. Each sample was then pipetted into multiple wells of a 96-well half-area, low-binding, clear-bottomed PEG coating plate (Corning 3881), at 80 μL per well.

For the seeded experiments, preformed fibrils were freshly prepared just before the experiment without sonication because it has been shown previously that sonicating the fibrils for 10 min in a sonicator bath to delump the fibrils has no effect on their capacity to accelerate secondary nucleation^15^. ThT experiments were set up just as above for 5 μM Aβ42 samples in 20 mM sodium phosphate buffer (pH 8) with 200 μM EDTA, and 20 μM ThT. The ThT fluorescence was monitored for 3.5 h to verify the formation of fibrils. Samples were then collected from the wells into low-binding tubes. The final concentration of fibrils, in monomer equivalents, was considered equal to the initial concentration of monomer. Fibrils were then added to freshly prepared monomer to reach a 0.5%, 1%, 5%, or 10% final concentration of seeds.

### AFM imaging

High-resolution and phase-controlled AFM was performed on positively-functionalized mica substrates. 10 μL of 0.5% (v/v) 3-aminopropyl-triethoxysilane (APTES, Sigma) in Milli-Q water was deposited onto freshly cleaved mica and incubated for 1 min. The substrate was rinsed three times with 1 mL of Milli-Q water and dried by a gentle stream of nitrogen gas. Finally, for each sample, an aliquot of 10 μL of the solution was deposited on the functionalized surface. The droplet was incubated for 5 min, then rinsed with 1 mL of Milli-Q water and dried under nitrogen gas. The preparation was carried out at room temperature. AFM maps were acquired using an NX10 AFM (Park Systems) operating in non-contact mode and equipped with a silicon tip (PPP-NCHR, 42 N/m) with a nominal radius <10 nm. Image flattening was performed by SPIP (Image Metrology) software.

### Fabrication of microfluidic spray devices

A two-step photolithographic process was used to fabricate the master used for casting microfluidic spray devices. In brief, a 25 μm thick structure was fabricated (3025, MicroChem) was spin-coated onto a silicon wafer. This, was then soft-baked for 15 min at 95 °C. An appropriate mask was placed onto the wafer, exposed under ultraviolet light to induce polymerization, and then post-baked at 95 °C. A second 50 μm thick layer (SU-8 3050, MicroChem) was then spin-coated onto the wafer and soft-baked for 15 min at 95 °C. A second mask was then aligned with respect to the structures formed from the first mask, and the same procedure was followed, i.e. exposure to UV light and post-baking for 15 min at 95 °C. Finally, the master was developed in propylene glycol methyl ether acetate (Sigma-Aldrich) to remove any photoresist which had not cross-linked.

A 1:10 ratio of PDMS curing agent to elastomer (SYLGARD 184, Dow Corning, Midland, MI) was used to fabricate microfluidic devices. The mixture was cured for 3 hours at 65 °C. The hardened PDMS was cut and peeled off the master. The two complementary PDMS chips are then activated with O2 plasma (Diener Electronic, Ebhausen, Germany) and put in contact with each other and aligned precisely such that the gas inlet intersects with the liquid inlet to form a 3D nozzle ^13^.

### Use of microfluidic spray devices

Prior to introduction of sample, each device was tested and washed with MilliQ water for 5 min. Sample was then loaded into 200 μL air-tight glass syringes (Hamilton) and driven into the spray device using a syringe pump (Harvard apparatus). Solutions containing sample were pumped into the device with a flow rate of 100 μl/h, while the nitrogen gas inlet pressure was maintained at 3 bar. Deposition was conducted for a maximum of 15 s at a distance of 4.5 cm to ensure coalescence did not occur. Samples were sprayed directly onto the FTIR prism with no further washing steps required before measurements.

### FTIR spectroscopy

Fibrils at 5 μM (monomer equivalent) were retrieved directly from an aggregation reaction and deposited onto the prism as described above. Measurements were performed on a Vertex 70 FTIR spectrometer (Bruker). Each spectrum was acquired with a scanner velocity of 20 kHz over 4000 to 400 cm^−1^ as an average of 200 scans using a *DiamondATR* unit and a deuterated lanthanum α-alanine-doped triglycine sulfate (DLaTGS) detector. New background spectra were acquired before each measurement. All spectra were analysed using OriginPro (Origin Labs). Spectra presented represent 3 individual spectra which were averaged and normalised. To determine the secondary structure composition of proteins, a second derivative analysis was performed. Spectra were first smoothed by applying a Savitzky-Golay filter.

