## Supplementary material for "Maturation-Dependent Changes in the Structure and Seeding Capacity of Aβ42 Amyloid Fibrils": Si Information

**Abstract:** Many proteins self-assemble to form amyloid fibrils with a cross-b sheet structure, a process which has implications in both human disease, such as neurodegenerative disorders, and in functional material development. Thus, the aggregation process has been widely studied, shedding light on the properties of fibrils and their intermediates. However, mature amyloid fibrils obtained at the plateau phase of the aggregation reaction are often treated as uniform, static structures, with little attention paid to the heterogeneous features which emerge over time. Here, we study the properties of mature amyloid-b fibrils as a function of time to challenge this long-held notion. We demonstrate that mature fibrils display a maturation process, with an increase in both fibril length and thickness, and a change in the cross-b sheet content. These changes affect the capability of the fibrils to act as templates for secondary nucleation. This has consequences for both our understanding of fibril maturation processes, as well as for the use of amyloid fibrils in drug discovery and materials science applications.

### 2. SI Figures

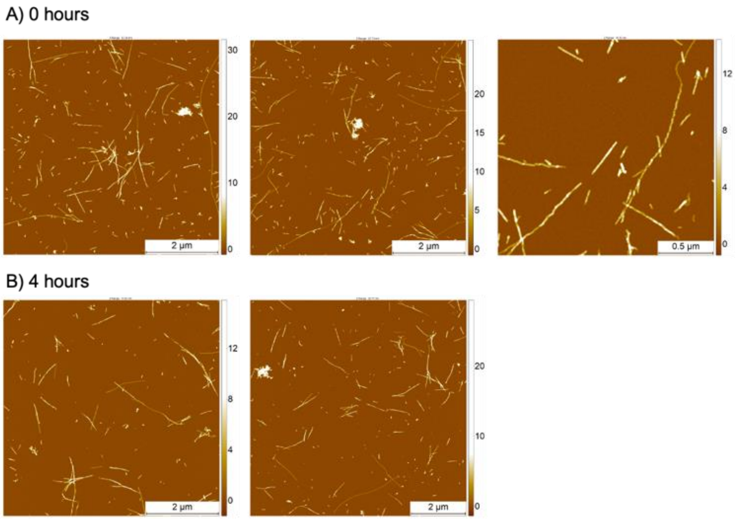


**Figure S1.** Example AFM images of samples acquired at a) 0 and b) 4 hour time points in the plateau phase.

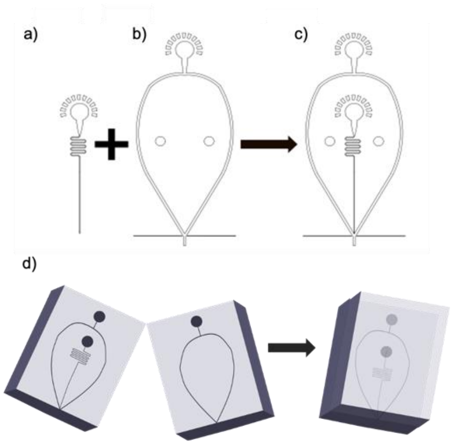


**Figure S2.** Schematic of the microfluidic device assembly. A 25 μm high liquid channel (a) and a 50 μm high gas channel (b) are combined in a two-step lithography process (c). Masters are used to fabricate two complementary PDMS slabs, which are carefully aligned and bonded to form the 3D spray nozzle device (d).

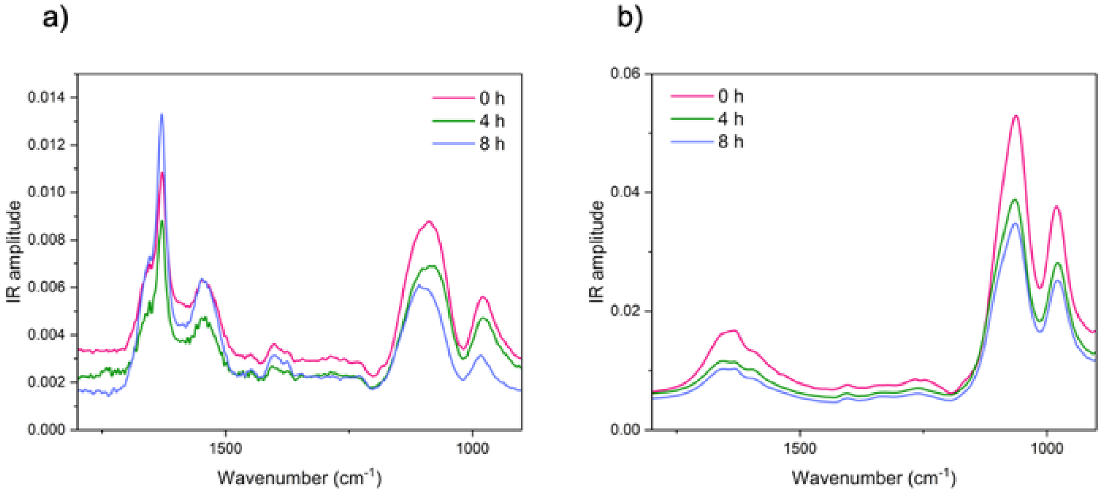


**Figure S3.** Non-normalised spectra of Aβ42 fibrils at 5μM in sodium phosphate buffer are shown for spray deposition (a) and manual deposition (b) to demonstrate the difference in spectroscopic sensitivity between sample preparation methods. Ultra-fast drying of picolitre-volume droplets from spraying results in reduced time for salt crystals to form. Salt crystallisation can be monitored by the peaks at ~1070 $cm^{-1}$. The intensity of the amide I peak is higher than that of the salt peak when deposited via spray. On the other hand, the large contribution of salt to the spectrum for manual deposition masks the protein amide I and II peaks, preventing secondary structure analysis.

### 4. Author Contributions

AM, SC and FSR performed the experiments. AM, SC and FSR analysed the data. AM and FSR wrote the original draft. All authors contributed to the final manuscript.
